# Structure-based self-supervised learning enables ultrafast prediction of stability changes upon mutation at the protein universe scale

**DOI:** 10.1101/2023.08.09.552725

**Authors:** Jinyuan Sun, Tong Zhu, Yinglu Cui, Bian Wu

## Abstract

Predicting free energy changes (ΔΔG) is of paramount significance in advancing our comprehension of protein evolution and holds profound implications for protein engineering and pharmaceutical development. Traditional methods, however, often suffer from limitations such as sluggish computational speed or heavy reliance on biased training datasets. These challenges are magnified when aiming for accurate ΔΔG prediction across the vast universe of protein sequences. In this study, we present Pythia, a self-supervised graph neural network tailored for zero-shot ΔΔG predictions. In comparative benchmarks with other self-supervised pre-training models and force field-based methods, Pythia outshines its contenders with superior correlations while operating with the fewest parameters, and exhibits a remarkable acceleration in computational speed, up to 10^5^-fold. The efficacy of Pythia is corroborated through its application in predicting thermostable mutations of limonene epoxide hydrolase (LEH) with significant higher experimental success rates. This efficiency propels the exploration of 26 million high-quality protein structures. Such a grand-scale application signifies a leap forward in our capacity to traverse the protein sequence space and potentially enrich our insights into the intricacies of protein genotype-phenotype relationships. We provided a web app at https://pythia.wulab.xyz for users to conveniently execute predictions. Keywords: self-supervised learning, protein mutation prediction, protein thermostability

## Introduction

Proteins, often described as the molecular workhorses of biology, perform an astonishing variety of biochemical duties integral to life^1,2^. Despite their vital roles, most natural proteins are marginally stable, with Gibbs free energy differences between the native and unfolded states as low as 5 kcal/mol^3^. This narrow stability margin renders them highly susceptible to perturbations from environmental changes or genetic mutations^4^. Even minuscule alterations, such as single point mutations, can tip this delicate balance and propel an active protein towards nonfunctional, misfolded, or aggregated forms. The implications of these changes stretch far and wide across health, disease pathology, pharmaceutical development, biotechnology, and our understanding of protein evolution^5^.

In this era, marked by efforts to transcend the limitations of natural protein repertoires, protein engineering has emerged as a promising avenue^6^. Protein sequences have been designed with enhanced stability, solubility, and tailored activities to fulfill the demand of industrial applications^2,7^. Computational tools have been meticulously crafted for protein design, following both model-based and data-driven methodologies^8^. As an important model-based methodology, the energy calculation allows for predictions of ΔΔG (difference of ΔG between the wildtype and mutant) upon amino acid substitution to identify stable mutations^9^. Several studies have leveraged well-developed energy functions such as Rosetta^10^, FoldX^11^, and ABACUS^12^, achieving good success in the design of thermostable enzymes^13^. However, they are still suboptimal because of the imbalanced parametrization of the energy function, and insufficient sampling of conformational space^9^. The incomprehensibly vast size of the hyper-dimensional sequence space is a continuing challenge for exploring protein sequence space^14^.

In light of this, recent advances in machine learning offer a promising solution. Models explicitly designed for ΔΔG mutation predictions predominantly employ a supervised regression framework. These models typically rely on carefully curated evolutionary features, such as BLOSUM62^15^ and probabilities in multiple sequence alignments (MSAs)^16^, or structural features including surface accessible area^17^, predicted H-bonds^18^, atomic charges^19^, Rosetta/FoldX modeled mutation structures^19^ and energy terms^20,21^. Although appealing for directly learning from experimental data and offering improved speed, supervised methods grapple with the scarcity of experimentally measured ΔΔG training data and biases present in these training datasets^22, 23, 24^. Such issues are pervasive in biology due to the laborious nature of wet-lab experiments^25^. In contrast to supervised learning, which is limited by labeled data, self-supervised learning (SSL) is able to learn from vast unlabeled data^26^. A particularly prevalent strategy within SSL is masked language modeling (MLM), which trains models to predict a masked or replaced token based on its surrounding context^27^. MLM has found widespread application across protein sequences^28^, MSAs^29^, and protein structures^30^, especially in the prediction of mutation fitness. For instance, ESM-1v^31^, trained with MLM on 150 million sequences from the UniRef90 database, achieved superior zero-shot fitness prediction on 41 deep mutation scanning experiment datasets with an averaged Spearman’s rho of 0.509. In addition to sequence-based pretraining, the ABACUS-R trained on a high quality subset of protein structure data, have shown success in *de novo* protein design^32^.

Guided by the general principles of SSL, we developed Pythia, a self-supervised model tailored for ΔΔG prediction of mutations based on protein structures. Constructed to decode intrinsic patterns among residues within given proteins, Pythia paves the way for precise anticipation of mutational impacts. This model operates independently of both evolutionary information and manually derived features from energy functions. Instead, it learns the stability directly from the protein structures. Upon evaluation with thousands of widely trusted experimental ΔΔG data, Pythia exhibited superior performance in both prediction correlation and computational speed, surpassing current state-of-the-art models. By spotlighting limonene epoxide hydrolase (LEH), we empirically displayed Pythia’s ability to identify a significantly higher proportion of effective thermostable mutations. Further, we highlighted Pythia’s potential for comprehensive exploration of the protein universe by calculating all single mutations from the high-quality predicted structures in the AlphaFold database^33^, totaling over 26 million predicted protein structures. Pythia’s source code is freely accessible at https://github.com/Wublab/pythia.

## Results

### 1. Model architecture and training of Pythia

In the past few decades, many methods have attempted to explore the correlation between the free energy landscape and the internal structure of proteins, but their accuracy seems to be limited by the approximations and assumptions inherent in the models. In this context, we hypothesized that the energy of a protein in its unfolded state is largely unaffected by mutations^15^, as there are virtually no stable specific interactions between the side chains of a protein in an unfolded state: ΔΔ*G* ∼ *G_MUT_* − *G_WT_*. Simultaneously, the probability of each amino acid represents the sum of the probabilities of all rotamer conformations of this amino acid over the entire rotamer space of all 20 amino acids. This probability can be transformed into a sum of each conformation and the energy corresponding to that conformation, i.e.,

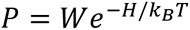

where each amino acid’s microstate (conformation) count and entropy are related as:

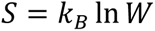

substituting into the above equation:

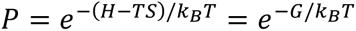

from this, we can derive that ΔΔG is proportional to the amino acid probability:

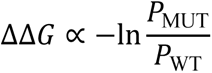

We therefore adopted a widely used graph neural network architecture extensively recognized for its successes in protein structure prediction^34^ and sequence design^35^. A protein local structure was transformed into a graph representation using a k-nearest neighbor graph (Figure 1A). Each amino acid served as a node in the graph, forming connections with its 32 closest amino acids based on the Euclidean distance of the C-alpha atom. The node features encompassed one-hot encoding for the amino acid type and the backbone dihedral angles (*φ*, *ψ*, and *ω*) encoded using the sine and cosine. To ensure SE(3)-invariance, the distances between the five backbone atoms were considered for the edge encoding, including C-alpha, C, N, O, and C-beta when available. In addition to the fundamental features, supplementary features were incorporated such as relative positional encoding of amino acids in the sequence and chain identity encoding. The chain identity encoding assigned a binary value of one if two amino acids belonged to the same chain, or zero if not (Figure 1B). The training objective aimed to predict the natural amino acid type of the central node based on the information from the nodes and edges (Figure 1C).

**Figure 1.**
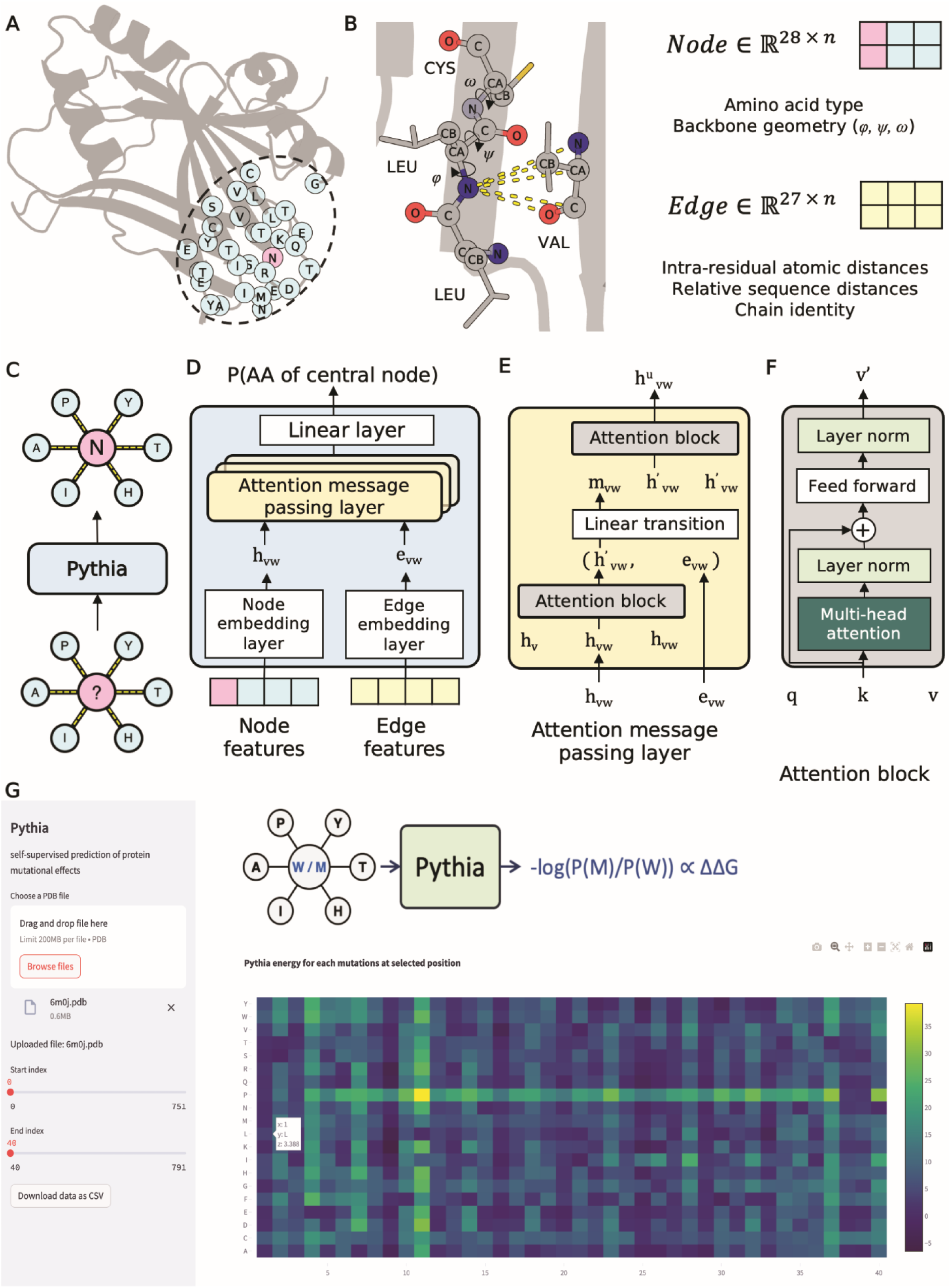
Overview of the Pythia model. (A) Pythia takes a protein’s local structure as input, represented as a k-NN graph derived from the Euclidean distance of C-alpha atoms. This local structure is abstracted into a graph composed of amino acids. (B) Node features encompass the type of each amino acid (including a MASK token), and three dihedral angles: *φ*, *ψ*, *ω*. Edge features include the distances between main chain atoms, sequence positions, and chain information. (C) The training task of Pythia is to predict the correct amino acid type of the central node. (D) The architecture of the Pythia model. The features of nodes and edges independently traverse the embedding layer and then enter the attention-based message passing neural network. The final output is the probabilities of the 20 amino acids. (E) Breakdown of the architecture of attention message passing layer. Within this layer, the information of nodes is first updated using the attention block. The embeddings of edge (*e_vw_*) are concatenated with the representation of nodes (*h’_vw_*) to get *m_vw_*. Subsequently, *m_vw_* and *h’_vw_* go through the attention module, resulting in the updated updated *h*^u^*_vw_*. (F) The structure of the attention block. (G) A visual snapshot of the Pythia webapp inerface.

Pythia employed the message passing neural network (MPNN) architecture^36^, specifically tailored with an attention-based message passing and read-out function (Figure 1D). By integrating attention mechanisms with MPNN, referred to as the Attention Message Passing Layer (AMPL), the model is extended to focus more on substructures critical to the desired interaction properties during the learning process. In each layer of the AMPL, the vertex representation was updated using an Attention block and concatenated with the edge representation to obtain the message representation (Figure 1E). The message representation then served as a query to further update the node representation through another Attention block (Figure 1F). The final model comprises three AMPLs, each processing a hidden dimension of 128.

Throughout the model training phase, we assessed several hyperparameters, including the masking ratio of the central nodes and the noise level of the backbone coordinates, as detailed in Table S1. To ensure both robustness and generalizability, we developed two separate models. One model was trained on pre-defined protein domains sourced from the CATH database^37^, whereas the other was nurtured on a non-redundant protein structure dataset built in this study by clustering high resolution bio-assemblies from the RCSB PDB database^38^. The final prediction of Pythia is made by using the averaged output of these two models. We provided a web app at https://pythia.wulab.xyz to run predictions (Figure 1G).

### 2. Benchmark evaluation of Pythia in ΔΔG prediction

Pythia was evaluated alongside a diverse set of pretrained protein models and three commonly utilized energy function methods on the S2648 dataset^39^, a standard training set for supervised machine learning models for predicting ΔΔG of mutations due to its high quality. Throughout this assessment, Pythia achieved a Spearman’s rho of 0.616 and Pearson’s r of 0.598 (Figure 2A), outperforming all models evaluated across six critical performance metrics (Spearman’s rho, Pearson’s r, accuracy, F1-score, AURCO, and AUPRC) (Figure 2B). It is noteworthy that all structure-based pretrained model showed higher correlation compared to sequence-based and MSA-based pretrained models, with the state-of-the-art model (ESM2-t33^40^) failing to achieve a correlation surpassing 0.4. This underscores the advantage of incorporating structural information, which provides inter-residue interaction information, as a more effective strategy for predicting the thermodynamic properties and mutation-induced effects of protein. Nonetheless, when compared with energy function-based methods, the structure-based models remain suboptimal, with Pythia standing out as a notable exception that achieved a higher correlation. Pythia demonstrated a superior ability to transfer to the single mutation prediction task, and for the first time, outperformed force field-based methods in pre-trained models. Remarkably, Pythia accomplishes this with only 1.3 million parameters, which is one-third of the second smallest model (Figure 2C).

**Figure 2.**
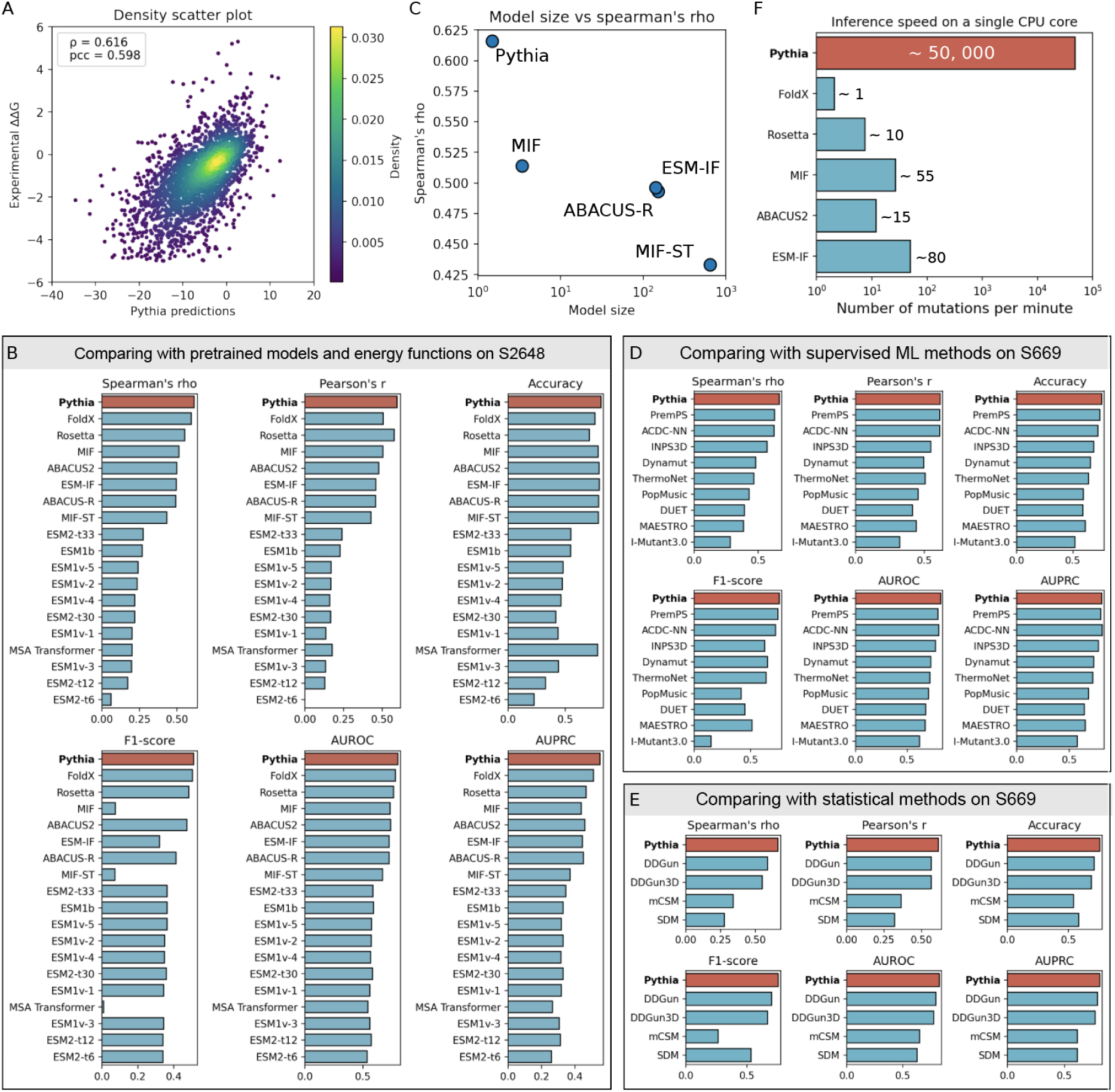
Evaluation of Pythia in predicting ΔΔG compared with current state-of-the-art methods. (A) The density scatter plot for Pythia’s predictions. (B) Parameters of the top five deep learning methods and their Spearman’s rho on S2648. (C) A comparison of the inference speed among the top six methods ranked by the Spearman’s rho. (D) Comparisons of Pythia against pretrained models and energy functions. The correlation of the predicted values with experimental ΔΔG is indicated by both Spearman’s rho and Pearson’s r, revealing the ranking and linear correlation. The metrics for classification tasks (Accuracy, F1-score, AUROC and AUPRC) aim to accurately categorize the stabilizing factor (ΔΔG_folding_ > 0) using the S2648 dataset. (E) Comparisons of Pythia against 9 supervised machine learning methods using the S669 dataset. (F) Comparisons of Pythia against 4 knowledge-based statistical methods using the S669 dataset. The top-performing method has been highlighted in red and the remaining methods are depicted in blue.

We further explore the performance of Pythia against supervised machine learning models. A direct comparison with machine learning-based predictors is challenging due to their diverse training datasets causing possible data leak and baises^41^. We therefore used a S669 dataset, which has not been used to train by supervised machine learning models and shares sequence identities less than 25% with S2648 and the varibench dataset^23^. As shown in Figures 2D and 2E, the prediction performances reach a Spearman’s rho ranging from 0.28 to 0.63 for supervised machine learning models and 0.28 to 0.59 for statistical methods. Pythia outperformed the methods evaluated in this study on the S669 dataset across all metrics, with a Spearman’s rho achieving 0.66. A major problem for supervsied ΔΔG predictors lies in their inability to ensure symerticity of direct and inversed mutations^23^. Comparing with these methods, Pythia dose not rely on any labeled ΔΔG data during the training stage and can produce perfect symertical between direct and inversed mutation with fixed protein backbone (Tabel S2).

Beyond the prediction accuracy, Pythia boasts an additional advantage of computational speed. While force field-based methods require local sidechain or even backbone structural conformation sampling to achieve more accurate prediction, they are constrained by their diminished computational speed, particularly when handling large-sequence-length proteins. Even when the backbone is fixed, force field-based methods can only manage to compute about 10 mutations per minute. Among the tested force field-based methods, FoldX, which is the best performing, is even slower. It manages an average of a single mutation per minute on a CPU core (E3-2678v3) due to its extensive sampling methodology, which involves multiple independent runs and averaging. When tasked with the computation of 380,741 mutations for 131 proteins in the S2648 dataset, Pythia requires only 20 seconds for initialization and computation on 24 CPU cores, approximating 0.5 million mutations per minute on a single core. This outperforms other methods by a factor ranging from 10 to 50 thousand (Figure 2E). By employing fewer local feature representations, the computational resource requirement is further optimized due to less preprocessing computation. Consequently, the computational demand for a single mutation within the neural network remains consistent regardless of an increase in protein length.

### 3. Evaluation of Pythia on a mega-scale dataset

Expanding the scope of our study, we applied predictions to a mega-scale dataset of approximately one million mutations distributed over 600 proteins including natural, redesigned and hallucinated domains^42^. The performance was evaluated on 258, 435 mutations within 181 well-characterized natural protein domains. The overall Spearman’s rho is 0.606 and the Pearson’s r is 0.623 (Figure 3A), while the AUROC reached 0.83 (Figure 3B) and AUPRC reached 0.76 (Figures 3C), consistent with the performance on S2648 and S669. Out of 181 tested natural domains, 127 domains (70%) have a Spearman’s rho higher than 0.6 (Figure 3D), a statistic suggestive of a relatively strong correlation^43^. This observation prompted us to further explore the domain-specific correlation. Unlike a global evaluation across all point mutations, assessing the correlation value for individual domains offers insights better suited for applications in protein engineering and mutation prediction. By analyzing each natural protein domain in this manner, we achieved a higher average Spearman’s rho of 0.620. An illustrative example emerged in our analysis of the structural domain of Ssl0352 protein from *Synechocystis sp.* PCC 6803 (PDB ID: 2JZ2). The correlation between scores predicted by Pythia and the ΔΔG values estimated through cDNA display proteolysis reached a notable Spearman’s rho of 0.778 (Figure 3E), indicating a remarkably strong relationship.

**Figure 3.**
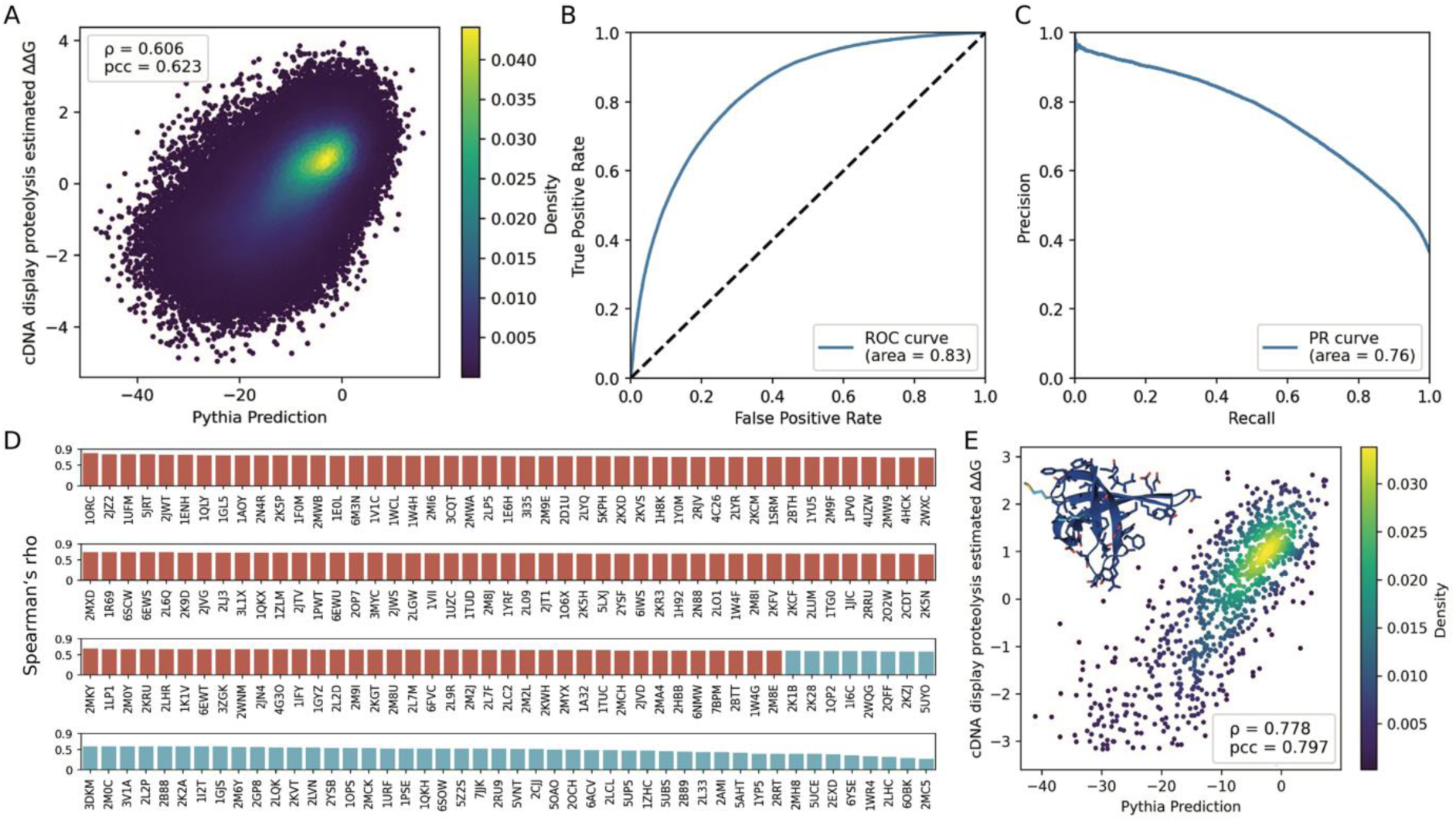
(A) Density scatter plot showcasing Pythia’s predictions on the mega scale dataset. (B) ROC curve illustrating Pythia’s ability to classify stabilizing mutations. (C) PR curve highlighting Pythia’s prediciton accuracy in classifying stabilizing mutations. (D) The correlation between Pythia’s predictions and the cDNA display proteolysis estimated ΔΔG is represented by Spearman’s rho across all 181 domains (E) Density scatter plot of Pythia’s predictions for the structural domain of Ssl0352 protein.

### 4. Identification of stabilizing mutations for a limonene epoxide hydrolase

Encouraged by the superior generalization in predicting ΔΔG, we experimentally verified Pythia’s predictions using the LEH from *Rhodococcus erythropolis* DCL14. This enzyme has been used widely in organic chemistry and has undergone extensive protein engineering, allowing for direct comparisons between different strategies^44^. We selected mutations with scores below −2 predicted by Pythia, prioritizing those with the lowest scores when multiple mutations were possible at a given position. This process led to a total of 36 single point mutations (Figure 4A, Table S3), among which 17 mutations resulted in an increase in the protein’s melting temperature (*T*_m_) (Figure 4B). Hybrid strategies employ visual inspections by molecular dynamic simulations (MDs) to filter out unreasonable candidates predicted by energy function methods such as FoldX, thus improving the median Δ*T*_m_ from −1.80 °C to - 0.15 °C. However, not only does this require a high level of technical expertise, but it also hinders the widespread adoption of such a strategy. By comparison, whereas hybrid strategies required knowledge-based mutation selection, Pythia improved the median Δ*T*_m_ to 0.90 °C of all expressed variants without any expert intervention (Figure 4C). Notably, the proportion of mutations with a *T*_m_ increase exceeding 1°C was significantly higher in Pythia’s predictions compared to energy function methods, even with MDs-based filtration (Figure 4D). Among these beneficial mutations, the P57A mutation, which is typically regarded as destabilizing in force field-based methods, exhibited the highest *T*_m_ increase of 8 °C. Moreover, only four of the 17 beneficial mutations had been previously reported, highlighting Pythia’s unique capability to identify stabilizing mutations that may have been overlooked by conventional methods. The laborious experimental evaluation of extensive mutation libraries often involves sifting through thousands of variants, with most random mutations proving neutral or detrimental to stability. Pythia’s aptitude to expand the landscape of beneficial mutations offers a valuable supplement to existing stability prediction tools, broadening the repertoire of beneficial mutation sequences accessible through computational predictions. In light of this, Pythia emerges as a promising foundational tool to bolster the advancement of hybrid strategies, such as FireProt^45^, FRESCO^44^ and GRAPE^13^, that integrate information from diverse complementary approaches to provide more options for the subsequent accumulation paths.

**Figure 4.**
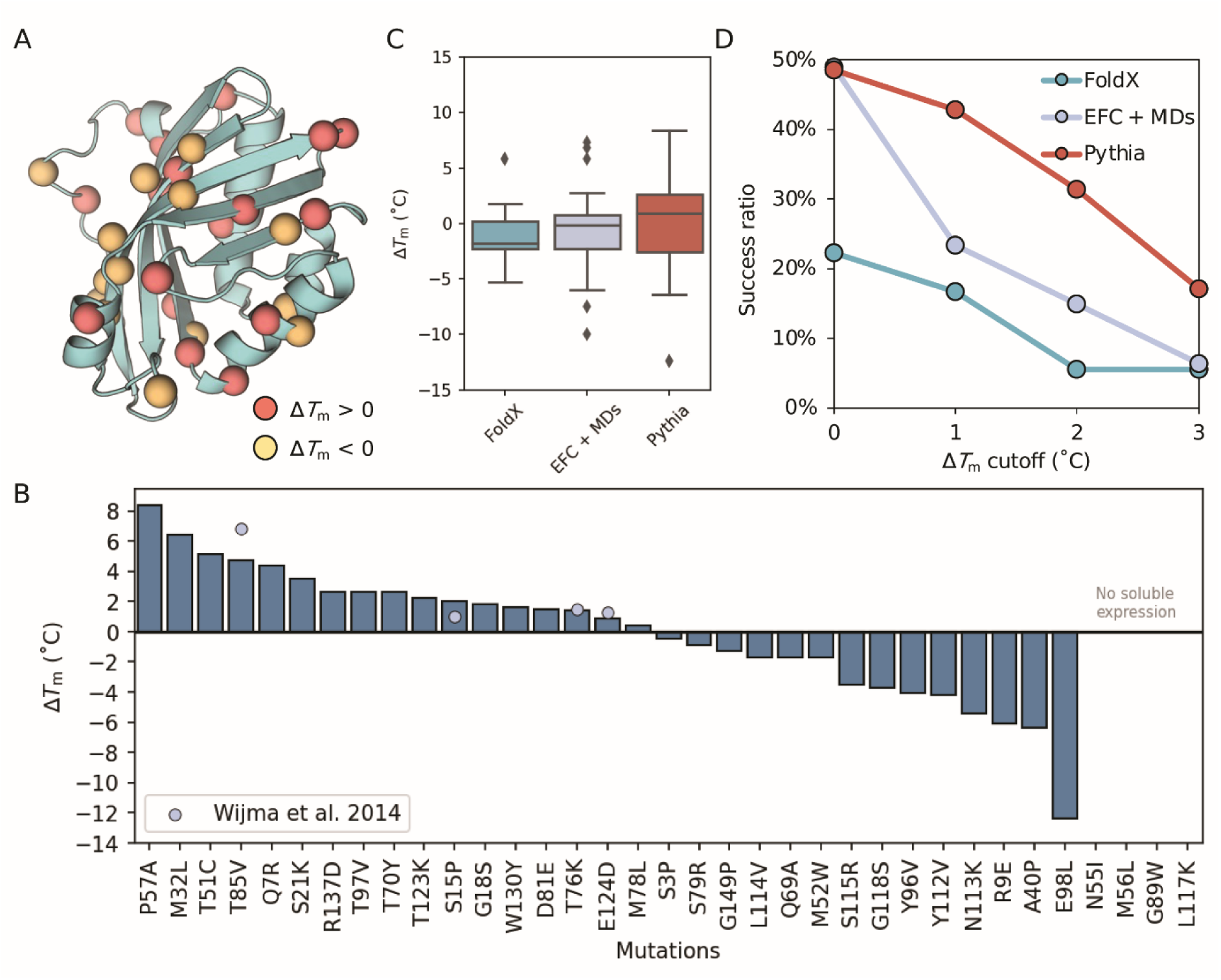
(A) The monomer structure of LEH is rendered in cartoon form with the C-alpha atom locations of mutations shown as spheres. Stabilizing mutations are represented with red spheres, while destabilizing mutations are shown in yellow. The visulazation of protein structures were prepared using PyMOL. (B) A bar plot representing the measured Δ*T*_m_ of variants characterized in this study. Bars accentuated with light blue dots present the stabilizing mutations as reported by Wijma et al.^44^ (C) A boxplot comparing three different mutation prediction strategies. The central line within each box represents the median values of Δ*T*_m_ for that strategy. The top and bottom boundaries of the box represent the first and third quartiles, respectively. The height of the box represents the interquartile range (IQR). Data points outside of the 1.5*IQR range are considered outliers and are plotted individually. (D) The success ratio of characterized mutations versus different Δ*T*_m_ cutoff values across three strategies. The blue curve depicts results using only FoldX. The grey curve represents the outcomes of energy function calculations supplemented with further MDs filtration (termed EFC + MDs), which corresponds to the single-point mutation prediction component within the FRESCO strategy. The red curve demonstrates results achieved using only Pythia, devoid of any other selection criteria.

### 5. Structural interpretability of Pythia

Since Pythia employs an attention mechanism, we can leverage the attention scores learned by the model to investigate whether it has indeed succeeded in capturing the complex interactions within proteins. We visualized the attention scores of functional residues in molecular graphs from two distinct categories in Figure 5. The first instance considers the π-π interactions of F52 and its neighboring residues within the GB1 domain (PDB ID: 1PGA). Four aromatic amino acids, Y3, F30, W43, and Y45, are located in the proximity to F52 and potentially form π-π interactions with F52 to stabilize the hydrophobic core of the domain (Figure 5A). Intriguingly, Pythia assigns higher attention scores to these four amino acids and the pivotal F52, indicating the model’s aptitude to discern the importance of these interactions relative to other neighboring residues (Figure 5B). Regarding pre- and post-mutation structures, we examined DuraPETase^13^, a more stable plastic-degrading enzyme engineered from *Is*PETase. Previous studies have suggested the synergistic effect of D186 with multiple stability-enhancing point mutations. We contrasted the attention scores allocated to the mutated residues surrounding D186 with their wild-type counterparts (Figure 5C). The results revealed that Pythia apportions higher attention scores to the mutated interactions, suggesting that Pythia is sensitive to the structural implications of mutations and can effectively capture the consequential relationships between mutated residues and their surroundings (Figures 5D, 5E).

**Figure 5.**
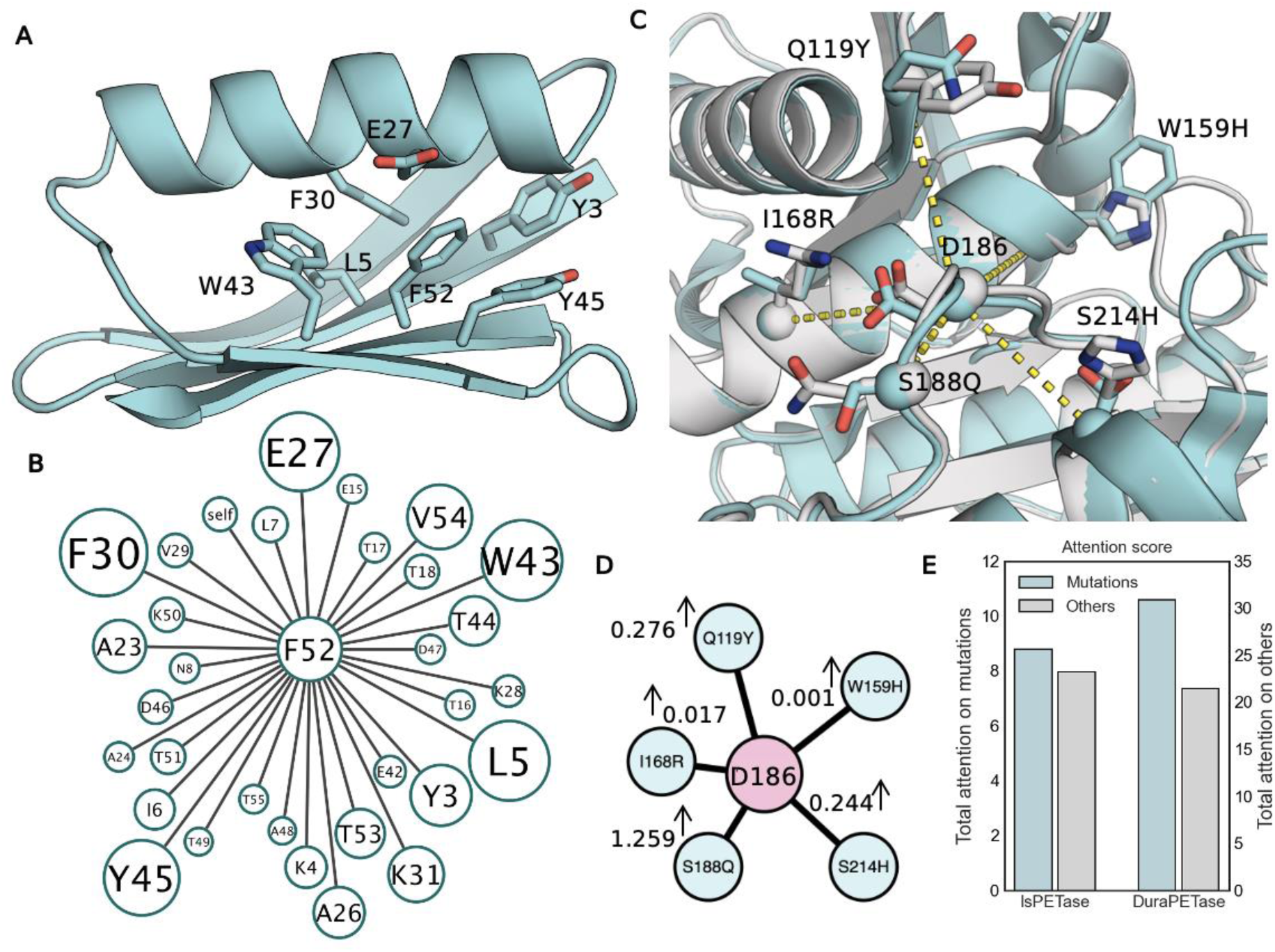
Interpretability of Pythia. The attention score can be interpreted as a measure of the impact of amino acids in the environment on the central amino acid distribution. (A) The π-π interactions of F52 with its neighboring residues in the GB1 domain (PDB ID: 1PGA). Possible π-π interactions are highlighted with blue dashed lines. (B) The k-NN graph of the F52 in the GB1, the weights are assigned based on the attention score in the final AMPL. Thicker and shorter egde indicate a higher attention score. (C) A comparison between *Is*PETase (PDB ID: 5XH3) with the DuraPETase (PDB ID: 6JKI). *Is*PETase is colored in green and the DuraPETase in white. The sidechain of mutation positions and the central node D186 is shown in sticks and their C-alphas are displayed as spheres. (D) The change in attention scores between mutated residues surrounding D186 with their wild-type counterparts. (E) The sum of total attention scores at the five mutated positions and positions that remain unchanged. The visualization of protein structures were prepared with PyMOL.

### 6. ΔΔG prediction at the protein universe scale

We further investigated the prediction speed of Pythia on three different levels: 1) proteome scale, 2) annotated proteins, and 3) the protein universe. For proteome-scale speed assessment, we used the proteome of *Escherichia coli* K-12 as a representative example (Figure 6A). Pythia swiftly predicted all 25,189,782 mutations for 4,214 structures (with an average C-alpha pLDDT > 70) in just 3 minutes using a single NVIDIA GeForce RTX 4090. Next, we expanded our effort to include all mutation predictions for the 134,276 structures in SwissProt with pLDDT score above 95. Impressively, this vast computational effort was finished in roughly 2 hours, scanning a total of 770,105,473 mutations. Lastly, we processed all possible mutations for 26 million high quality AlphaFold2 structures. A computational challenge of such magnitude would be virtually insurmountable for traditional tools like FoldX, which would demand more than 30 years to handle this on 1,000 CPUs. In stark contrast, Pythia completed the entire computation in 3 days using a machine equipped with 8 NVIDIA GeForce RTX 4090 GPUs. This clearly showcases Pythia’s immense computational efficiency for large-scale mutation prediction.

**Figure 6.**
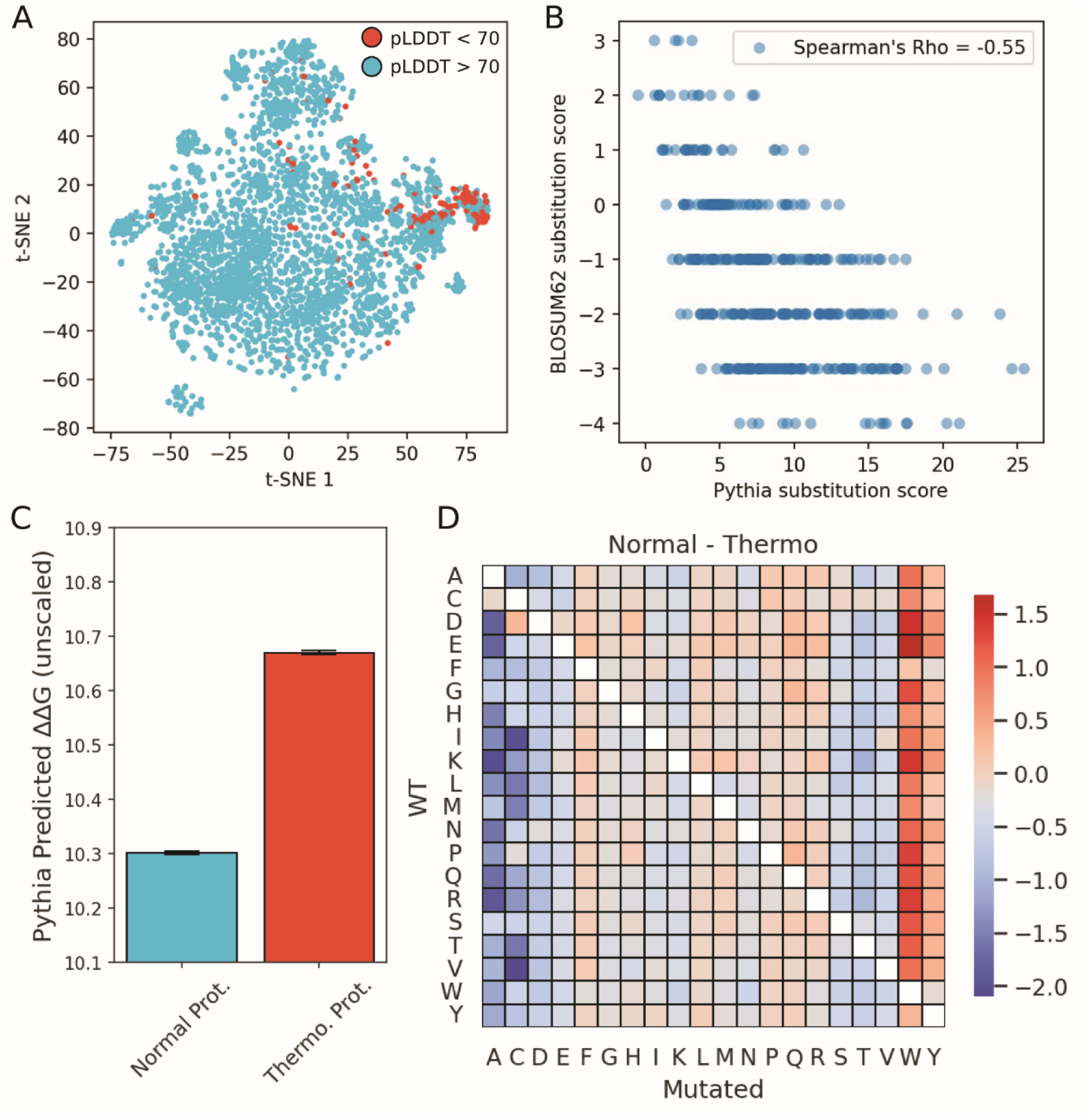
(A) Visualization of the protein space with a t-SNE embedding of the *E.coli* K-12 proteome. Blue dots represent proteins with averaged C-alpha pLDDTs equal to or higher than 70, while red dots represent proteins with averaged C-alpha pLDDTs less than 70. (B) Scatter plot comparing amino acid substitution scores of Pythia and BLOSUM62. (C) Bar plot depicting the averages of all mutations. Mutations in thermotolerant proteins exhibit significantly higher Pythia score (P value = 0.0 in Mann-Whitney U test) compared to mutations in random sampled proteins (most likely to be non-thermostable). (D) Heatmap illustrating energy differences caused by various mutation types. The 380 mutation types are colored according to the averaged energy difference.

In our preliminary analysis of millions of mutations derived from a uniform distribution, we found that the average scores of amino acid substitutions correlated with the substitution scores in the BLOSUM62 matrix (Figure 6B). This suggests that the driving force behind the evolution might be tied to the fundamental thermodynamics of amino acid substitutes. Furthermore, we identified a significantly higher averaged mutation score in the thermophilic proteins than non-thermophilic proteins (with a p-value of 0.0 from the Mann-Whitney U test, Figure 6C). Although the difference is slight, it indicates that sourcing stabilizing mutations from a thermostable scaffold might be more challenging, inferring a more limited sequence space for thermophilic proteins. Drawing upon a comprehensive dataset of mutational variations, we undertook an analysis into the role of residue type in influencing protein stability by comparing thermophilic and non-thermophilic proteins. A clear pattern emerged in the predicted mutations that smaller substituents (A or C) might be generally favorable. In contrast, substitutions with aromatic rings (F, Y, and W) seem to be disadvantageous in thermophilic proteins (Figure 6D). By utilizing the advantages of self-supervised learning, our investigation into large-scale protein mutations unveiled intricate details often missed in isolated protein mutation studies.

## Discussion

The prediction of free energy changes (ΔΔG) upon mutation plays an integral role in deciphering the impacts of genetic variations on protein function and stability. Such knowledge is pivotal for advancements in fields like personalized medicine and protein engineering. Given the dearth of labeled ΔΔG data required for deep learning, we present Pythia, an efficient approach designed for zero-shot predictions that harness the power of self-supervised learning. The unique architecture of Pythia permits integration of both sequence and structural data, thereby improving the prediction accuracy by accounting for the multidimensional influences on protein stability. Pythia’s self-supervised training component allows it to ascertain unbiased amino acid distributions based on local structures. Furthermore, its attention weights yield valuable biological insights, making patterns of interactions interpretable and bolstering the model’s explainability.

Comparative evaluations with other self-supervised pretraining models underscore Pythia’s superior Spearman correlation and Pearson correlation, achieved with the fewest number of parameters. When pitted against traditional energy calculation methods, Pythia not only surpassed these alternatives but also exhibited a remarkable computational speed increase of up to 10^5^-fold. The efficacy of Pythia is further evidenced by the prediction of 1 million experimentally characterized mutations and its successful application in thermostable mutation prediction for LEH. Such computational efficiency holds particular promise for large-scale, high-throughput investigations across expansive protein datasets.

Several ways are available for enhancing Pythia’s performance and broadening its range of applications. Evolutionary information has been recognized as a crucial feature in both supervised machine learning models and pre-trained models. The integration of evolutionary information, such as Position-Specific Scoring Matrices, into the training target could potentially improve Pythia’s performance while maintaining its computational efficiency^47^. A current constraint of Pythia is its reliance on predicted structures as a starting point. However, with 152 billion genetic variations predicted in the present study, it seems feasible to align pre-trained protein language models with the probabilities predicted by Pythia. Such incorporation may facilitate more accurate and unbiased sequence-based ΔΔG prediction of mutations. Additionally, the emergence of large-scale biological prediction datasets, as exemplified by AlphaFold protein structure database, represents a significant advancement in the field of computational biology. Fusion mutation information with structure datasets may unlock the potentials of these predictive datasets, advancing our understanding of the complexity of biological phenomena on a previously inconceivable scale.

## Methods

### 1. Details of feature and training of Pythia

The task is to predict the best fitted amino acid type for a given local chemical environment of a protein. The local chemical environment is defined by the *k*-nearest neighbor of the central residue. As the residue type is determined by interactions of its neighbors, therefore the network should recognize both the residue types and their relative geometries. To extract the geometric features, atomic-pairwise distances of five backbone atoms were used. Besides the geometric features, relative sequence position encoding and chain identity encoding were combined to get the SE(3)-invariant edge attributes *X* ∈ ℝ^27^. The node attributes *X* ∈ ℝ^28^ combined amino acid type embedding (*X* ∈ ℝ^22^, 20 canonical amino acids, X for all uncanonical amino acids and the mask token) together with sin and cos encoded backbone torsional angles (psi, phi, and omega). The node attributes are also SE(3)-invariant. For model input, the token of central amino acid was replaced to the mask token at a given probability or randomly substituted to the left 21 tokens for the reset. The model was trained with a cross entropy loss and optimized using Adam optimizer with a learning rate of 3e-4.

The non-redundant data sets of protein domains from the CATH database clustered at 40% identity was used for model training and evaluation. 31 domains were removed because they had an amino acid type labeled as UNK and 270 domains shorter than 32 amino acids were removed. We randomly split the domains into 29,800 for training, 200 for validation, and 1,584 for testing. The non-redundant dataset of protein biological assemblies was downloaded from the RCSB PDB database and yielded 27,999 protein assemblies meet the following criteria:

1. Resolution < 2.5 Å and Refinement R Factors (R Work) < 0.25
2. Sequence length between 32 to 10000
3. Structure solved using X-ray diffraction
4. Protein polymer entities grouped by 50% sequence identity
5. Released before 2020-05-01

We downloaded representatives and randomly selected 400 biological assemblies for model validation.

During model training, geometric features were perturbed with Gaussian noise levels of 0, 0.02, 0.1, 0.2, 0.3, 0.4, and 0.5. The perturbations were only added to the atomic distances. During inference, no noise was added.

On the CATH domain dataset, the model with only 1.3 million parameters thus can be trained overnight (about 9 hours) on a personal laptop equipped with an RTX 3070 graphic processing unit (GPU). For efficiency, we trained the model at Bscc Cloud computation cluster with 4 RTX 3090 GPUs within one hour. The model was implemented in Python using PyTorch and Lightning packages.

### 2. Zero-shot ΔΔG prediction

Pythia is trained to predict the probabilities of each amino acid at a certain position given a protein structure context.

The masked marginal score is computed more efficiently as it predicts probabilities of all amino acids at the given position from a mask noise as the following:

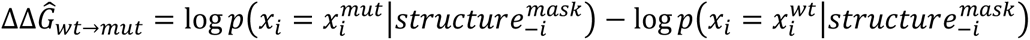

For all unsupervised predictions, we utilized the masked marginal score, which is fast to compute because only one forward pass is required for a specific mutation’s position, and by employing the masked marginal score, symmetry is assured.

The only exception is ABACUS-R. Since it lacks a specific token for masking at the output layer, we adopted the ΔΔlogit method, as originally proposed in its publication.

### 3. ΔΔG predicted with energy functions

For FoldX, the input structure was amended using the RepairPDB module to sample and repack side-chains using the FoldX force field. Once the protein structure was repaired, mutations were introduced using the *BuildModel* module. To strike a balance between sufficient sampling and time cost, the *--numberOfRuns* was set to 5 by default. The final prediction was made by averaging the 5 scores.

For Rosetta, the structure was initially relaxed using the *fastrelax* protocol. The relaxed structures were then processed using the *ddg_monomer* application to calculate ΔΔG, employing the fastest protocol row1^47^: -ddg::weight_file soft_rep_design -ddg::iterations 1 - ddg::local_opt_only true -ddg::min_cst false -ddg::mean false -ddg::min true -ddg::sc_min_only false-ddg::opt_radius 0.1.

For ABACUS, the default setting was applied using the *singleMutation* to compute energy terms of a single mutation, and the ΔΔG was calculated as the sum of all energy terms.

### 4. Mega scale experimental dataset cleaning up

We evaluated the performance of Pythia using a recently published large-scale folding stability experiment^42^. Our analysis included variants with single amino acid substitutions that have well-defined experimental ΔΔG values. Mutations were classified into four categories: class 1: variants in natural protein domains with well-characterized ΔΔG, class 2: all variants for natural protein domains, class 3: variants derived from mutated natural protein domains, class 4: variants in *de novo* designed protein domains. Only the results of class 1 were included in the main text. This categorization was primarily due to the observation that structure prediction algorithms often perform poorly in predicting the structure of mutants and are not robust in predicting *de novo* designed proteins. These shortcomings can significantly impact the prediction results.

### 5. Mutation prediction on billion scale predicted structure

We filtered the AlphaFold database and selected structures with an overall pLDDT score greater than 95 and length between 32 and 1280. This yielded a total of 26,648,233 structures.

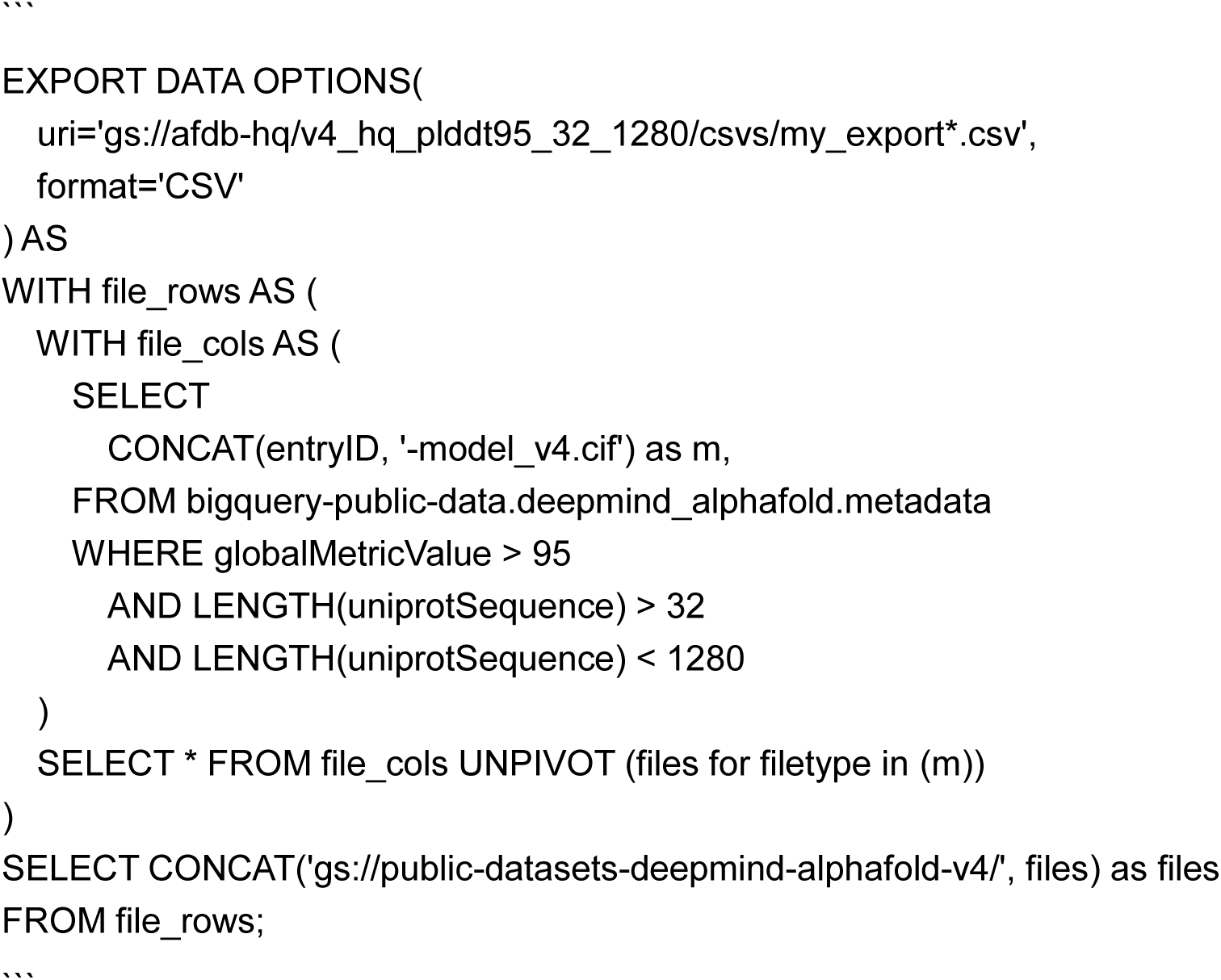

We downloaded all mutations to a local machine, resulting in approximately 5 TB of data, which was divided into 1667 subsets. Each subset contained approximately 16,000 structures, roughly the size of a eukaryotic proteome, larger than a typical prokaryotic proteome. The computation was conducted by parallel processing 6 structures on a RTX 4090 GPU. It took about 13 minutes to finish computation of a subset. The prediction throughput was approximately 7 million mutations per minute per card.

We also tested the speed on a very modest machine, with 2 E5-2678v3 CPUs and 128G DDR3 memory. It required about 30 minutes to finish computation of a subset.

### 6. Webapp and API

The webapp was built with streamlit, can be easily accessed at https://pythia.wulab.xyz. The API was built with the FastAPI package, with and endpoint of /scan/, user can upload a PDB file and get predictions returned in plain text.

curl -X POST “http://pythia.wulab.xyz/scan/?output_format=2col” -F “file=@.pdb” >

results.txt

### 7. Metrics

We used the Spearman’s rho and Pearson’s r to measure the correlation between predicted and experimental ΔΔG.

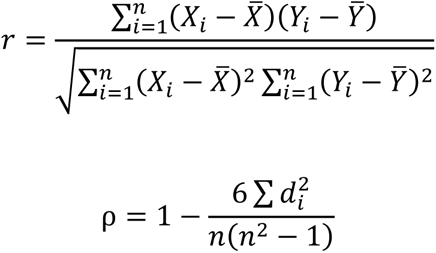

For classification, we used accuracy, F1-score, area under the receiver operating characteristic curve (AUROC) and the area under the precision-recall curve (AUPRC) to systemically compare the classification performance of all tested methods.

### 8. Expression, purification and characterization of the LEH mutants

The genes of the wild-type LEH and its mutants were synthesized and cloned into the vector pET-21a(+) by General Biosystems (Anhui, China) with sequence optimization for expression in *E. coli*. To prepare the samples for determining the apparent melting temperature, *E. coli* BL21(DE3) cells carrying the expression vector pET21a-LEH were cultured in 50 mL of LB broth containing 50 μg/mL of ampicillin at 37°C, 180 rpm until optical density at 600 nm (OD_600_) reached approximately 0.8. At that point, protein expression was induced by the addition of 1 mM isopropyl thio-β-D-galactoside (IPTG) followed by incubation at 20°C, 180 rpm for 16 h. The cells were collected by centrifugation (8,000 g, 10 min, 4°C) and lysed by sonication on ice in buffer A (containing 50 mM KH_2_PO_4_, 200 mM NaCl and 20 mM imidazole, pH 7.5). After removal of precipitates via centrifugation (14,000 g, 60 min, 4°C) and filtration (0.22 μm filter, Millex), the cell extract was obtained, then loaded onto a 5 mL HisTrap HP column (GE Healthcare). The enzyme was collected by elution with buffer B (containing 50 mM KH_2_PO_4_, 200 mM NaCl and 300 mM imidazole, pH 7.5), then imidazole and NaCl were removed using a HiPrep 26/10 desalting column (GE Healthcare) with buffer C (containing 50 mM KH_2_PO_4_, pH 7.5). The purified enzyme was concentrated to 0.5 mg/mL via Amicon filtration (10 kDa, Millipore), then stored at −20°C until use.

The apparent melting temperatures (T_m_) of the LEH mutants were determined by using Prometheus NT.48 system (NanoTemper Technologies, Munich, Germany). A sample of 10 µl of protein solution (0.5 mg/mL in 50 mM KH_2_PO_4_, pH 7.5) was heated from 25 to 85°C at a heating rate of 1.0°C/min. Fluorescence changes at 330 nm and 350 nm were monitored, respectively.

