## Supplementary material for "Structure-based self-supervised learning enables ultrafast prediction of stability changes upon mutation at the protein universe scale": SI

Table of Contents

Notes ........................................................................................................................2

Additional tables ........................................................................................................3

**Notes**

1. Hyperparameter of Pythia

Before training Pythia, we conducted an evaluation to assess the influence of various mask ratios, replacement ratios, and noise levels on the model using main-chain coordinates. The evaluation involved training the model for up to 40 epochs using structural domain data sourced from CATH 4.3. The data was divided into training, validation, and testing sets in an 8:1:1 ratio. Model selection was based on accuracy on the validation set, and subsequent comparison involved examining accuracy on the testing set as well as the Spearman correlation coefficient on the ΔΔG dataset (S2648).

Our findings indicated that the mask ratio had a relatively minor impact on the model's performance. However, as the noise level on the coordinates increased, the correlation coefficient for predicting ΔΔG gradually improved, eventually reaching a plateau at around a 0.4 to 0.5 Å noise level (Table S1). Consequently, we opted for a mask ratio of 0.85 and a noise level of 0.5 Å for the final model configuration. Subsequently, we trained two models, Pythia-C and Pythia-P, on CATH and protein biological assemblies data, respectively.

### Additional tables

Table S1. Experiments on choosing the noise level and mask ratio

| Mask vs Substitute | Training noise (Å) | Test accuracy (%) | Test Loss | Test perplexity | Spearman's rho on S2648 |
|---|---|---|---|---|---|
| 0.85 vs 0.15 | 0.00 | 0.56 | 1.34 | 3.84 | 0.53 |
| | 0.02 | 0.55 | 1.40 | 4.06 | 0.54 |
| | 0.10 | 0.52 | 1.51 | 4.50 | 0.57 |
| | 0.20 | 0.50 | 1.56 | 4.78 | 0.57 |
| | 0.30 | 0.49 | 1.61 | 4.99 | 0.57 |
| | 0.40 | 0.48 | 1.64 | 5.13 | 0.59 |
| | 0.50 | 0.48 | 1.66 | 5.25 | 0.59 |
| 0.60 vs 0.40 | 0.00 | 0.56 | 1.34 | 3.84 | 0.52 |
| | 0.02 | 0.55 | 1.40 | 4.05 | 0.54 |
| | 0.10 | 0.52 | 1.50 | 4.46 | 0.57 |
| | 0.20 | 0.50 | 1.56 | 4.77 | 0.57 |
| | 0.30 | 0.49 | 1.60 | 4.97 | 0.57 |
| | 0.40 | 0.48 | 1.63 | 5.12 | 0.59 |
| | 0.50 | 0.48 | 1.65 | 5.21 | 0.58 |

Table S2. Comparison of Pythia with existing methods on S669

| Method | Spearman's rho | | | Antisymmetry | |
|---|---|---|---|---|---|
| | Total | Direct | Inverse | $r_{d-i}$ | $\langle\delta\rangle$ |
| **Pythia** | **0.66** | **0.46** | **0.46** | **-1** | **0** |
| PremPS | 0.63 | 0.42 | 0.42 | -0.82 | 0.38 |
| ACDC-NN | 0.63 | 0.45 | 0.44 | -0.99 | 0.09 |
| INPS3D | 0.57 | 0.44 | 0.36 | -0.46 | 1.02 |
| Dynamut | 0.48 | 0.37 | 0.36 | -0.55 | 0.62 |
| ThermoNet | 0.46 | 0.37 | 0.34 | -0.83 | 0.31 |
| PopMusic | 0.43 | 0.41 | 0.22 | -0.23 | 1.42 |
| DUET | 0.39 | 0.42 | 0.23 | -0.07 | 1.48 |
| MAESTRO | 0.39 | 0.46 | 0.19 | -0.17 | 1.32 |
| I-Mutant3.0 | 0.28 | 0.35 | 0.15 | -0.01 | 1.62 |
| DDGun | 0.59 | 0.43 | 0.41 | -0.98 | 0.12 |
| DDGun3D | 0.55 | 0.43 | 0.41 | -0.97 | 0.18 |
| mCSM | 0.34 | 0.36 | 0.21 | -0.02 | 1.75 |
| SDM | 0.28 | 0.39 | 0.14 | -0.42 | 0.97 |

Table S3. Expression and measured $T_m$ of 35 variants and the wildtype of LEH

| Mutation | Soluble expression | $T_m$ (˚C) | $\Delta T_m$ (˚C) | Pythia Score |
|---|---|---|---|---|
| T51C | Yes | 56.5 | 5.1 | -4.175 |
| T85V | Yes | 56.1 | 4.7 | -7.363 |
| Q7R | Yes | 55.8 | 4.4 | -4.47 |
| S21K | Yes | 54.9 | 3.5 | -3.512 |
| T123K | Yes | 53.6 | 2.2 | -4.415 |
| S15P | Yes | 53.4 | 2.0 | -7.331 |
| G18S | Yes | 53.2 | 1.8 | -2.536 |
| T76K | Yes | 52.8 | 1.4 | -2.11 |
| E124D | Yes | 52.3 | 0.9 | -2.385 |
| S3P | Yes | 50.9 | -0.5 | -10.391 |
| M52W | Yes | 49.7 | -1.7 | -8.492 |
| Q69A | Yes | 49.7 | -1.7 | -3.421 |
| Y96V | Yes | 47.3 | -4.1 | -4.171 |
| R9E | Yes | 45.3 | -6.1 | -2.913 |
| A40P | Yes | 45.0 | -6.4 | -3.674 |
| E98L | Yes | 39.0 | -12.4 | -2.179 |
| G89W | No | - | - | -4.791 |
| P57A | Yes | 59.8 | 8.4 | -3.585 |
| M32L | Yes | 57.8 | 6.4 | -5.633 |
| R137D | Yes | 54.0 | 2.6 | -2.579 |
| T70Y | Yes | 54.0 | 2.6 | -4.63 |
| T97V | Yes | 54.0 | 2.6 | -3.57 |
| W130Y | Yes | 53.0 | 1.6 | -2.691 |
| D81E | Yes | 52.9 | 1.5 | -2.108 |
| M78L | Yes | 51.8 | 0.4 | -4.489 |
| S79R | Yes | 50.5 | -0.9 | -2.889 |
| G149P | Yes | 50.1 | -1.3 | -5.628 |
| L114V | Yes | 49.7 | -1.7 | -4.251 |
| S115R | Yes | 47.9 | -3.5 | -3.833 |
| G118S | Yes | 47.7 | -3.7 | -2.024 |
| Y112V | Yes | 47.2 | -4.2 | -5.834 |

| Mutation | Soluble expression | $T_m$ (˚C) | $\Delta T_m$ (˚C) | Pythia Score |
|---|---|---|---|---|
| N113K | Yes | 46.0 | -5.4 | -2.593 |
| L117K | No | - | - | -2.793 |
| N55I | No | - | - | -8.086 |
| M56L | No | - | - | -2.345 |
| Wild type | Yes | 51.4 | 0.0 | - |
